# Meningeal vascular Aβ deposition associates with cerebral hypoperfusion and compensatory collateral remodeling

**DOI:** 10.1101/2025.02.05.635937

**Authors:** Alexandra M. Kaloss, Jack L. Browning, Jiangtao Li, Yuhang Pan, Sachi Watsen, Harald Sontheimer, Michelle H. Theus, Michelle L. Olsen

**Affiliations:** Department of Biomedical Sciences and Pathobiology, Virginia Tech, Blacksburg, VA, 24061; School of Neuroscience, Virginia Tech, Blacksburg, VA, 24061; Genetics, Bioinformatics and Computational Biology, Virginia Tech, Blacksburg, VA, 24061; Department of Neuroscience, University of Virginia, Charlottesville, VA, 22901

**Keywords:** CAA, Aβ, cerebral blood flow, pial collaterals, arteriogenesis, meninges

## Abstract

**Background:** Global reductions in cerebral blood flow (CBF) are among the earliest and most consistent abnormalities observed in Alzheimer’s disease (AD), preceding both cortical plaque formation and cognitive decline. While the pial arterial network—a critical supplier of intracortical perfusion—has been overlooked in this context, it may play a pivotal role in early vascular pathology. Here, we report extensive cerebral amyloid angiopathy (CAA) within the pial artery and arteriole network in the J20 (PDGF-APPSw, Ind) mouse model of AD.

**Methods:** Using premortem delivery of Methoxy-XO4 to label Aβ, and arterial vascular labeling, we assessed Aβ burden on the pial artery/arteriole network and cerebral blood flow in aged male and female WT and J20 AD mice.

**Results:** We show that 12-month-old J20 mice exhibit significant Aβ deposition across major leptomeningeal arteries (ACA, MCA) and pial collaterals, with ∼40% vessel coverage in males and ∼20% in females—substantially exceeding Aβ levels in cortical or hippocampal vessels. This vascular Aβ burden was accompanied by compensatory enlargement and increased tortuosity of pial collateral vessels. Yet, despite this apparent remodeling, CBF was reduced by ∼15% in J20 mice, and this decline was significantly associated with leptomeningeal CAA burden.

**Conclusions:** This is the first study to comprehensively characterize meningeal arterial Aβ accumulation in a preclinical model of vascular AD, mirroring recent observations in early-stage human disease. Our findings implicate meningeal CAA as a potential driver of early CBF disruption and suggest that pial collateral remodeling may reflect a compensatory response to vascular insufficiency. Moreover, we identify robust sex differences in CAA burden, paralleling sex-specific patterns of parenchymal Aβ pathology in humans. These results highlight the leptomeningeal vasculature as a novel and understudied locus for early AD pathology and a potential therapeutic target to preserve cerebrovascular integrity.

## Background

Alzheimer disease (AD) is an irreversible, neurodegenerative disease affecting over 6 million Americans; a number that is expected to increase to 13.8 million by 2060(1). Over 95% of individuals living with AD have idiopathic disease, yet amyloid beta (Aβ) accumulation and intracellular inclusions of hyperphosphorylated Tau protein represent pathological disease hallmarks across all individuals. Three rare single gene variants are known to cause AD, each involved in amyloid protein processing and breakdown. In AD, Aβ accumulates in the brain forming parenchymal plaques within and along blood vessel walls as cerebral amyloid angiopathy (CAA). These findings have driven decades of research aimed at understanding the precise nature between Aβ accumulation and central nervous system (CNS) pathology in AD. AD disease diagnosis typically occurs late in disease progression, and to date, no effective interventional strategies for the treatment of AD exist. It is widely believed that early intervention during the prodromal stage is critical for developing effective AD therapeutics.

Neuroimaging studies have revealed that reduced global cerebral blood flow (CBF) represents one of the earliest pathologies identified in the brains of individuals with AD(2–5). Studies performed in individuals with advanced AD, demonstrated lateralization of visuospatial and language deficits correlated with asymmetric reduced blood flow velocity and glucose metabolism, leading to the notion that reduced CBF was a consequence of reduced metabolic demand in atrophied brain tissues(2–5). However, prospective patient imaging studies and blood flow measurements in individuals with mild cognitive impairment (MCI) indicate reduced CBF occurs early in AD progression, prior to significant neurological decline(2, 4, 6, 7). Reduced CBF has received increasing attention as an early pathological mechanism in AD, which may drive reduced cortical tissue oxygenation(8) and glucose metabolism(8, 9) in individuals during the early stages of the disease. Importantly, reduced CBF and reduced glucose utilization are recapitulated in preclinical models of AD, particularly in models which present with high Aβ load(10–12). The underlying mechanism for reduced CBF is not well understood. However, several factors, including age, hypertension, high cholesterol, diabetes, cardiac disease, and obesity are associated with brain hypoperfusion and enhanced vascular risk in AD(13). Reduced parenchymal CBF may result from lower macrovascular blood flow in the intracranial arteries, including the middle cerebral artery (MCA), identified in individuals diagnosed with AD(14) and those with mild cognitive impairment(15). Results from the Alzheimer Disease Neuroimaging Initiative indicate reduced CBF correlates with global Aβ load, particularly in the early stages of AD(2).

As it relates to CAA, a recent meta-analysis indicates moderate to severe CAA is observed in 48% of individuals with AD(16). CAA is associated with failure of Aβ clearance, as well as pathological changes to the integrity of the blood vessel wall, inflammation, and intracortical circulatory disturbances(17). Aβ accumulates on small and medium intracortical vessels(17). Additionally, Aβ accumulation has been demonstrated on some meningeal arteries, and leptomeningeal anastomoses(18, 19). To date, however, little is known regarding the extent of Aβ accumulation across the meningeal vascular network and its relationship to intracortical blood flow observed in patients with AD and recapitulated in AD mouse models.

Here, we use the J20 mouse model to characterize the level and impact of CAA presence in the leptomeningeal surface on CBF. These mice develop robust parenchymal plaques and CAA by 6 months of age, outlining their utility in understanding the relationship between CAA and CBF.

Using Methoxy-XO4 to identify vascular amyloid, combined with arterial-specific labeling approaches we identified meningeal arterial vascular amyloid was significantly higher than intracerebral vascular accumulation, with total pial arterial vascular Aβ accumulation approaching 40% of the entire meningeal vascular network in male J20 mice at one year of age, 20% for female J20 mice. Additionally, pial arterial CAA was associated with increased diameter of pial collateral vessels, which has been shown to improve blood flow in instances of cerebrovascular obstruction(20). Despite the observed increase in collateral diameter, indicative of blood flow redistribution, overall cortical hemisphere perfusion, as measured by laser speckle contrast imaging, was significantly decreased in AD mice relative to age-matched WT littermates. Our findings demonstrate significant Aβ burden on the meningeal arterial network and associated reduced CBF. Further, we identified significant enlargement of pial collateral vessels indicating a redistribution of CBF during disease progression.

## Methods

### Animals

The mouse strain used for this research project, male B6J.Cg-*Zbtb20^Tg(PDGFB-APPSwInd)20Lms^*/Mmjax, RRID:MMRRC_034836-JAX, was obtained from the Mutant Mouse Resource and Research Center (MMRRC) at The Jackson Laboratory, an NIH-funded strain repository, and was donated to the MMRRC by Lennart Mucke, Ph.D., Gladstone Institute of Neurological Disease ((Cat # 034836-JAX, MMRRC stock #34836)(21). This common mouse model of AD, herein called J20 mice, expresses mutant human amyloid protein precursor (APP) with the Swedish (K670N/M671L) and the Indiana (V717F) mutations under the mouse PDGF promoter. In a subset of experiments, APPSwFlLon,PSEN1*M146L*L286V)6799Vas/Mmjax, common name 5xFAD mice (MMRC strain #034848-JAX) were utilized. These mice express human APP with the Swedish (K670N/M671L), Florida (I716V), and London (V717I) Familial Alzheimer’s Disease (FAD) mutations and human Presenilin-1 with two FAD mutations, M146L and L286V. Both transgenes are expressed under the mouse Thy1 promoter. All mutant mice demonstrate progressive age-related Aβ deposition.(21, 22) All studies were approved by the Institutional Animal Care and Use Committee at Virginia Tech (IACUC #21-079, 24-037). Animals were housed in an AAALAC approved facility with a 12 h light-dark cycle; food and water *ad libitum,* and provided paper huts for environmental enrichment. Both strains of hemizygous males were bred to wild-type (WT) C57/B6 females in house. Genotypes of offspring were confirmed by PCR of DNA isolated from tail snips. Mutant and WT animals of each strain were co-housed following weaning until indicated time points. During the duration of the experiments, mice were frequently monitored by vivarium staff for health concerns. If a mouse showed neurological signs (seizures, paralysis, abnormal body carriage) after recovery from anesthesia it was immediately euthanized. In general, if any (surgical or non-surgical) mouse showed decreased activity, hunched, ruffled fur, it was immediately euthanized. In total, 12 male and 11 female J20 mice, 12 male and 11 female WT J20 mice, 4 male and 5 female 5X FAD, and 4 male and 5 female 5XFAD WT mice ranging from 10-12 months of age weighing between 20-60 grams were used in this study.

### Laser Speckle Surface Blood Flow Imaging

Blood flow monitoring was performed through a thinned skull using a laser speckle contrast imaging system v5.0 (RFSL III Laser Speckle Imaging System, RWD, China). Briefly, the mice were anesthetized with 2.5% isoflurane-100% oxygen, and the skin was prepared by hair removal and disinfection. A separate set of experiments were conducted using ketamine (10 mg/mL)/xylazine (1 mg/mL) as the anesthetic agent **(Supplementary Fig. 8)** to assess differences in anesthetic choice. All mice received buprenorphine (Ethiqa XR at 3.25 mg/kg) for analgesia during induction and were closely monitored during recovery and healing for signs of discomfort or pain. A midline incision was made, and the skull was thinned as previously described using a bone drill with carbide bur and then maintained moisture using a continuous application of warmed saline(23). The mice were placed in a stereotaxic adapter and positioned under the LSCI with an operating distance of 15cm. Temperature was monitored throughout preparation and maintained between 37 ± 2°C during blood flow imaging. Perfusion was measured over 60 seconds using the HD temporal algorithm at 2.5X zoom. Perfusion units were recorded in a standardized region of interest and averaged for each hemisphere. A pseudo color threshold range was set from 25-525 for all representative images. Only mice with intact hemispheres in both sides were used for laterization analysis. The experimenter was blinded to genotype during all stages of data collection. Animals were randomly selected for LSCI procedures across different times of day to ensure that sampling was not biased by potential confounding effects related to the sleep/wake cycle or circadian rhythms. One WT and one J20 mouse were excluded from the analysis because skull penetration occurred during the LSCI surgery, introducing the risk of altered cerebral blood flow and unreliable measurements.

### Methoxy-XO4 Labeling

Here we utilized systemic administration of Methoxy-XO4 (4,4’-[(2-methoxy-1,4-phenylene) di- (1*E*)-2,1-ethenediyl] bisphenol, Tocris cat# 4920) to visualize cerebrovascular amyloid and parenchymal plaques. Methoxy-XO4, is a fluorescent amyloid β probe which has been shown to readily cross the BBB, utilizing a carbon-11 labeled Methoxy-XO4 approach(24). Once bound, Methoxy-XO4 remains bound to plaques (over 90 days) and coupled with *in vivo* imaging can be repeatedly administered to measure plaque growth(25, 26).Methoxy-XO4 can be utilized for postmortem plaque and cerebrovascular amyloid assessment, where it has been shown to label plaques throughout the brain and co-localize with thivoflavin-S, Crenezumab, as well as other commercially available anti-Aβ antibodies(24, 26–28). Methoxy-XO4 was used to label vascular Aβ as described. Methoxy-XO4 was solubilized in DMSO (final concentration 100 uM, 30 uL), and added to an equal volume (30 uL) of Kolliphor EL (Sigma cat # C5135). This was further diluted to 1 mL in 1X phosphate buffered saline (PBS), to create a working solution. Each animal was injected intraperitoneally with a final volume of 2uL Methoxy-XO4 solution (final concentration Methoxy-XO4 10 mg/kg) per gram of body weight, 24 hours prior to cardiac perfusion.

### Vessel Painting

To label the pial arteriole/collateral network we applied a vessel painting approach we have described previously(29, 30). Briefly, twenty-four hours post-Methoxy-XO4 injections to identify vascular Aβ, J20, 5x-FAD, or control mice were injected with heparin (2000 units/kg), and sodium nitroprusside (SNP, 0.75 mg/kg). Five minutes post heparin injection, animals were euthanized with isoflurane, perfused with 10 ml of 1X PBS containing 20 units/ml heparin before switching to 10 ml 1,1’-Dioctadecyl-3,3,3’,3’-Tetramethylindocarbocyanine Perchlorate (Dil Stain, 0.01 mg/ml, Invitrogen, Cat # D282) diluted in 4% sucrose–PBS-heparin mixture at a flow rate of 2 ml/min. Dil is an orange/red lipophilic membrane stain (emission 549 nm) that labels vascular endothelial cells. Dil labeling was followed by ice-cold 4% paraformaldehyde (PFA), pH 7.4 to fix the tissue.

### Vessel painting image analysis

High-resolution tiled images at 4X magnification were taken of the surface of vessel painted brains using a Nikon C2 confocal (TI2-C-TC-I, Tokyo, Japan). Scaled mosaic images were imported into ImageJ-FIJI version 2.14.0 (NIH) to assess collateral number, tortuosity, and diameter, as described. Total number of inter-tree collaterals were identified between the MCA, ACA, and PCA artery branches. Collateral vessels were identified based on their characteristic wide or non-physiologic angle of insertion into the arterial trees(31), and tortuosity, combined with their unique positioning between the arterial trees of major arteries, directly connecting two arteries, as previously published(28, 32, 33) and seen in **Supplemental Fig. 1**. Here non-physiological angles refer to angles associated with a deviation from normal blood flow, where the wider bifurcation angle of blood vessels affects both shear stress and laminar flow, leading to more turbulent flow and a higher likelihood of pathologies(34–36). The diameter of collateral vessels was measured using the straight-line tool at 3 random points spanning the vessel and averaged across the brain. For collateral number, the average was taken between both hemispheres and reported for each niche. Total identified collaterals per hemisphere ranged from 3-18 for MCA-ACA collaterals, from 2-10 for MCA-PCA collaterals, and 0-3 for ACA-PCA collaterals. Collateral length and span were measured to determine collateral tortuosity. Length was measured with the straight-line tool by drawing a straight line from the start to the end point of the collateral. Span was measured with the freehand line tool by tracing the collateral from the start to end point. Tortuosity index was calculated as a ratio of the span:length of each collateral. For all imaging and collateral analysis, the experimenter was blinded to the condition.

### Fast 3D Clearing and Immunolabeling

Twenty-four hours after Methoxy-XO4 injections, animals were anesthetized with 500 mg/kg ketamine and 20 mg/kg xylazine via intraperitoneal injection and transcardially perfused with 20 ml cold phosphate-buffered saline (PBS), followed by 30 ml of 4% PFA in 1X PBS, pH 7.4. Brains were post-fixed overnight at 4°C in 4% PFA and then rinsed 5-6 times with PBS before immunostaining and clearing. In some cases, vessel painted brains were cleared. We utilized a fast 3D clearing protocol previously described(37) which includes a preclearing step, antibody staining and fast clearing.

*Preclearing*: Briefly, PFA fixed brains were incubated in several solutions with increasing concentrations of tetrahydrofuran solutions (THF, Millipore-Sigma cat #186562) to dehydrate the tissue. Fixed brains were rotated end over end in the following solutions: 50% THF, 50% water with 0.1% (v/v) triethylamine (pH 9.0) (Millipore-Sigma T0886) for 20 minutes at 4°C, then 70% THF, 30% water with 0.3% (v/v) triethylamine (pH 9.0) for 20 minutes at 4°C, then transferred to 90% THF, 10% water with 0.6% (v/v) triethylamine (pH 9.0) for 2 hours at 4°C. Rehydration was then achieved by rotating brains for 20 min each in 70% THF, 30% water with 0.3% (v/v) triethylamine (pH 9.0) followed by 50% THF, 50% water with 0.1% (v/v) triethylamine (pH 9.0) at 4°C. Samples are then washed with ddH20 five times for 10 min each wash at 4°C while protected from light. Brains were then immersed in a Fast 3D clearing solution (Histodenz (Millipore-Sigma D2158, final concentration 1.46 M), Diatrizoic Acid (Millipore-Sigma D9268, final concentration 24.43 mM), N-Methyl-D-Glucamine (Millipore-Sigma M2004, final concentration 128 mM), Sodium Azide (Millipore-Sigma S2002, 0.02% w/v) at 37°C until cleared.

*Immunolabeling:* Pretreated samples were incubated in PBS/0.2% Triton X-100 (Millipore-Sigma T9284)/20% DMSO (Millipore-Sigma D5879)/0.3 M glycine (Millipore-Sigma G8898) at 37°C 12 hours, then blocked in PBS/0.2% Triton X-100/10% DMSO/6% Teleostean gelatin l (FishGel, Millipore-Sigma G7041) at 37°C for 24 hours. Samples were washed in PBS/0.2% Tween-20 (Bio-rad 1610781) (PBST) for 1 hr twice, then incubated in primary antibody), goat anti CD31 (R&D Systems, AF3628, 1:300), goat anti Podocalyxin (R&D Systems, AF1556, 1:1500) dilutions in PBST/5% DMSO/3% FishGel 37°C for 3 days. Samples were then washed in PBST for 1 day, then incubated in secondary antibody donkey anti goat 488 (Thermo Scientific A-21202, 1:1000), dilutions in PBST/3% FishGel at 37°C for 3 days. Samples were finally washed in PBST for 2 days before clearing and imaging.

*Fast 3D Clearing:* After antibody incubation, brains were again rotated end over end at 4°C, protected from light, in successive increasing and then decreasing THF solutions to dehydrate and hydrate the tissue as described above with increased times: 1 hr for 50% and 70% THF solutions and 12-16 hours for 90% THF solutions. Finally, samples were washed with ddH20 4-5 times for 10 min each. Brains were then transferred into Fast 3D Clearing solution (3-5 ml for a single mouse brain) and incubated overnight at 37°C rotated end over end. Samples were always submerged completely in the solution. After overnight incubation or once samples were completely transparent, the samples were stored at 4°C protected from light.

### Imaging of cleared tissue and Image Analysis

Cleared tissue was imaged using a Nikon A1 confocal microscope (A1-SHS, Tokyo, Japan) using a 10X objective and 3.75µm step size, utilizing the z-intensity correction function at 512x512 resolution. For quantification of pial, cortical, and hippocampal Aβ load, Imaris (v9.8.2) 3D reconstructions of blood vessels, whole tissue, and Aβ plaques were created using 10x z-stack stitched images of vessel painted, CD31/podocalyxin and Methoxy-XO4 labeled brains. Blood vessels and Aβ plaques were reconstructed using the automated local threshold calculation in Imaris for unbiased quantification to account for background subtraction, with +/-50au manual adjustment frame, and a smoothing factor of 2.5µm. The volume ratio of Methoxy-XO4 to blood vessels was also recorded to determine percentage of vascular and parenchymal plaques. Here the volume ratio is defined as the percent of the vasculature with spatial overlap to Methoxy-XO4. Total Methoxy-XO4 coverage of blood vessels in the pial surface was compared to approaches previously employed quantifying surface coverage using a binary analysis of a compressed image in FIJI-ImageJ(18, 38–40). No statistical differences were observed between the two approaches (**Supplemental Fig. 2**). For the current study, we chose the Imaris analysis as it enabled 3D rendering.

### Assessment of Methoxy-XO4 and 6e10 Co-labeling

Coronal hippocampal and cortical brain sections (100 µm) from six J20 brains (2 slices each region from 5 male and 1 female) mice were collected to test the accuracy and penetration of Methoxy-XO4 labeling. Premortem Methoxy-XO4 labeling was co-labeled with postmortem 6e10 Aβ antibody, a commonly used antibody for labeling Aβ plaques(41–45), antibody using immunohistochemistry. Following a 5-minute wash in 1XPBS to remove cryoprotectant, each slice was blocked for 1 hour in 2% FishGel (Millipore-Sigma G7041) + 0.2% TritonX-100. Slices were incubated in a mouse 6e10 (1:500, BioLegend, cat #: 803001) and rabbit CoraLite 488 GFAP antibody (1:500, ProteinTech, cat #: CL488-16825) diluted in 1-part blocking buffer:3-part 1X PBS overnight at 4° C. Slices were washed three times in 1X PBS for 5 minutes each, and incubated with secondary antibody cocktail solution in diluted blocking buffer containing goat anti mouse 546 (1:500, Invitrogen, Cat #: A11003) for 1 hour at room temperature. Slices were washed three times in 1X PBS for 5 minutes each, and mounted onto a glass slide (Fisherbrand, cat #: 1255015) with mounting medium (Invitrogen, cat #: P36930) and subsequently imaged using a Nikon A1 confocal microscope with 0.5 µm step size using a 20X objective. Following imaging, 3D surface reconstructions were created in Imaris and the average number and size Methoxy-XO4 and 6e10 labeled plaques were recorded. Automated analysis in Imaris was used to assess co-labeled Methoxy-XO4 and 6e10 labeled plaques. This analysis was repeated using manual Image J with compressed Z-stack images. Plaques were characterized as bright, diffuse, spherical deposits of 100 voxels^3^ or larger.

*Leptomeningeal Imaris 3D (Oxford Instruments) reconstructions* of major arteries, distal arterioles, collaterals, and amyloid-β plaques were created using 4x z-stack, stitched images of vessel painted and Methoxy-XO4 labeled brains. Major arteries, distal arterioles, collaterals, and amyloid-β plaques were computed via absolute intensity, using a threshold value computed by creating a 3D region of interest around an area of the z-stack with no visible signal, in which the maximum intensity value +/- 100 of blood vessel painting and Methoxy-XO4 fluorescence was used as the minimum threshold value. Major arteries, distal arterials, collaterals and amyloid-β utilized a smoothing factor of 4.2µm. After surface reconstruction, arteries, distal arterioles and collaterals were identified and separated through manual clipping. Arteries, distal arterioles and collaterals were then labeled as Medial Cerebral Artery (MCA), Anterior Cerebral Artery (ACA), MCA Distal Arterioles, ACA Distal Arterioles, or MCA-ACA Collaterals. After major arteries, distal arterials, and collaterals were sorted, the percent coverage of amyloid-β on major arteries, distal arterials, and collaterals was recorded. For Imaris 3D analysis, the experimenter was blinded to the sex and genotype of the animal.

*Statistical Analysis*: Data was graphed using GraphPad Prism, version 9.5.1 (GraphPad Software Inc., San Diego, CA). Appropriate sample size was determined using a power analysis for a two-way ANOVA, assuming a power of 0.83 at an alpha level of 0.05, to detect significant differences in collateral vessel diameter between groups. This analysis estimated that n=10 mice per group would be required to achieve sufficient statistical power. Prior to statistical testing, a ROUT (Q=5%) outlier test was conducted, this resulted in 1 female J20 mouse being excluded from the study. For comparison of two experimental groups, a student’s two tailed t test was used. Paired t tests were used for data sets comparing lateralization. Multiple comparisons were performed using one-way ANOVA and repeated measures followed by *post hoc* Sidak test or a RM One-Way ANOVA and repeated measures followed by a *post hoc* Tukey test for matched comparisons. Changes were considered significant if p<0.05 and mean values were reported with standard error of mean (SEM).

## Results

Decades of research have consistently recognized reduced global CBF in individuals with AD. Early studies utilizing non-invasive transcranial Doppler ultrasound in AD patients relative to controls were used to demonstrate decreased blood flow velocity bilaterally in the MCAs which positively correlated with asymmetric cognitive impairment and regional cerebral glucose metabolism as measured by [^18^F]FDG-PET (5, 46). These studies led to the notion that reduced blood flow velocity in the MCA may result from reduced regional metabolic demands. However, this idea was challenged when a large, population-based, prospective study associated reduced CBF with incipient dementia and suggested cerebral hypoperfusion precedes and possibly contributes to clinical dementia(3). Now demonstrated across the spectrum of disease severity, in hundreds of studies, reduced CBF may represent an early AD disease biomarker(4, 7, 47) and reviewed in(48). Neuroimaging studies in patient cohorts indicate reduced CBF correlates with global Aβ load, particularly in early stages of AD(2), yet mechanisms underlying reduced CBF are unclear, and the contribution of meningeal CAA is not well characterized.

To address this gap, we utilized i.p. delivery of Methoxy-XO4 which has been shown to label CAA and parenchymal plaques(24, 27, 49) in J20 mice in comparison to WT littermates at one year of age(37). Here, we concurrently employed a select pial artery/arteriole tracing technique called *vessel painting* (VP) (19, 29), which enables detailed morphological analysis of the pial arterial and collateral network in aged J20 and WT mice. IMARIS 3D reconstructions of confocal imaged brains were generated for analysis (**Fig. 1A-B**). Examination of the pial surface Aβ coverage showed significant sex differences but no lateralization, with male mice having a significantly elevated CAA compared to females (**Fig. 1D**). This is the first evidence of sex differences and extensive pial arterial vasculature amyloid load in any murine model of AD. In the male J20 mice, there were no significant differences in Aβ coverage across the MCA, ACA, or interconnecting collaterals, indicating an equal distribution across the meningeal surface (**Fig 1C**). Female mice on the other hand, exhibited higher CAA in the ACA branch compared to the pial collateral vessels connecting the MCA-ACA branches (**Fig. 1C**).

**Figure 1.**
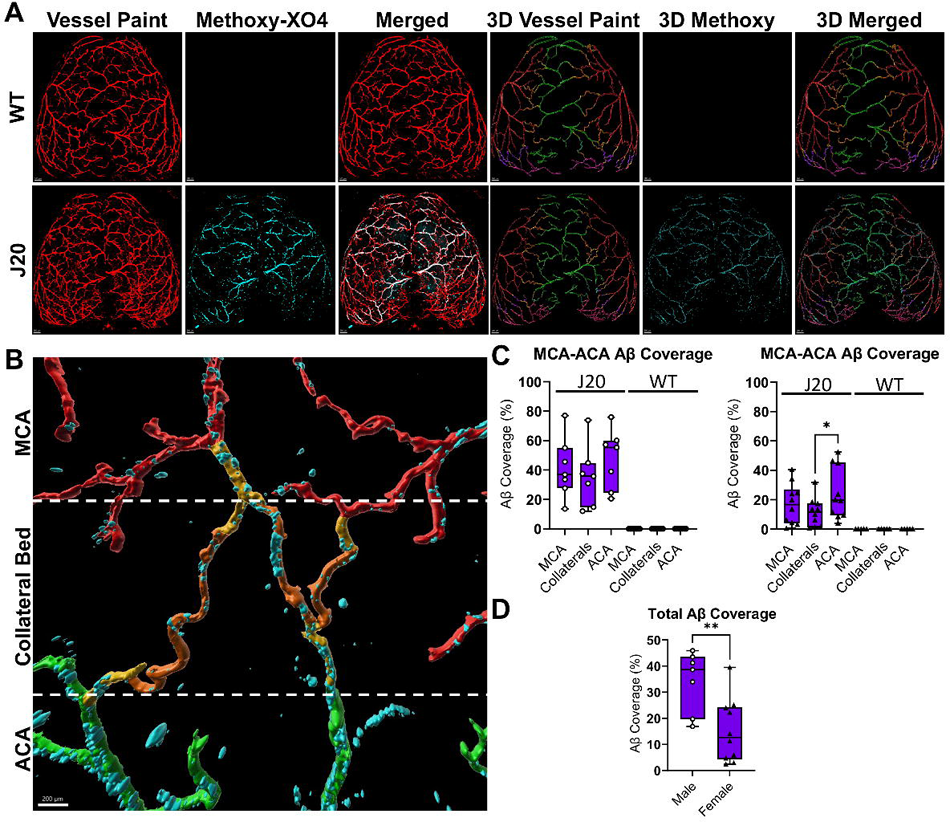
Leptomeningeal Aβ coverage in the 12 month old male and female WT and J20 mice. (**A**) Representative images of vessel paint (red) and Methoxy-XO4 (cyan) of the pial surface of 12-month-old male WT and male J20 mice brains and corresponding 3D Imaris reconstructions (scale bar = 500µm) (**B**) Representative zoomed image of MCA-ACA collateral bed in J20 animal (Anterior Cerebral Artery (ACA) = green, collaterals = orange, Medial Cerebral Artery (MCA) = red, distal arterials = gold, scale bar = 200µm) (**C**) % Aβ coverage of the MCA, MCA-ACA Collaterals, and ACA with significance detected by RM One-Way ANOVA test (top) or Ordinary One-Way ANOVA (Bottom) with Tukey’s multiple comparison test. (**D**). Comparison of % Aβ on all blood vessels in both hemispheres between male and female J20 mice with significance detected by unpaired t-test. n=7 Male J20, 10 Female J20, 7 Male WT, and 5 Female WT. * = p-value < .05. Circles = males; Triangles = Females

To determine if meningeal CAA and vascular changes in the J20 mouse line were strain-specific, 5xFAD mice were evaluated. IMARIS analysis of the meningeal surface showed Aβ accumulation on the main MCA and ACA, as well as the pial collateral vessels. Here we observed significant CAA, with levels similar to that observed for J20 female mice (**Supplemental Fig. 3**). Notably, unlike J20, 5xFAD mice displayed substantial Aβ deposition that was not vascular-associated and appeared throughout the meninges.

Brains also underwent Fast 3D clearing(37) and Imaris 3D reconstructions to quantify total pial surface, cortical and hippocampal Aβ load (**Fig. 2**). As demonstrated in the representative images, we observed significantly more vascular Aβ accumulation in superior axial brain sections at 1000µm thickness containing the pial cortical surface relative to cortical and hippocampal axial tissue sections of similar thickness in male J20 mice (**Fig. 2A**). These data indicate an enrichment of total Aβ load in the pial surface relative to parenchymal tissue, thus we next quantified total Aβ load in the pial surface, cortex, and hippocampus (**Fig. 2C)**. Our data indicates CAA is significantly elevated on pial arterial vasculature, whereas parenchymal Aβ was primarily identified as plaques, rather than CAA in 12-month-old J20 mice (**Fig. 2B, D**).

**Figure 2.**
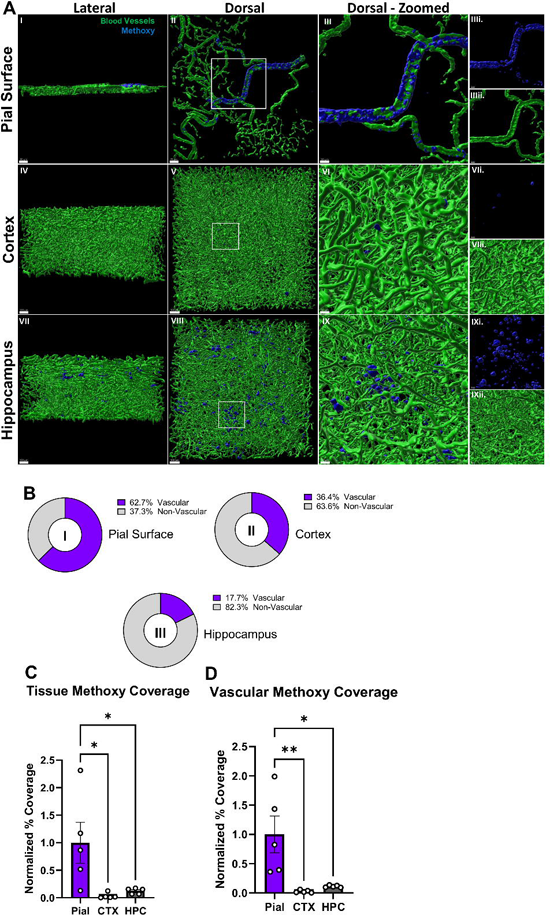
Vascular and nonvascular evaluation of amyloid-beta in 12mo J20 mice. (**A**) 3D Reconstructions of pial, cortical, or hippocampal blood vessels (green), and pial, cortical, or hippocampal Methoxy-XO4 (blue) from lateral (left) dorsal (middle) or zoomed dorsal (right) and zoomed dorsal individual channels views (individual channels denoted i. or ii.). Blood vessels of the pial surface were labeled with vessel paint and blood vessels of the cortex and hippocampus were labeled using Goat CD31 and Goat Podocalyxin primary antibodies in conjunction with Donkey anti Goat 488 secondary. (**B**) Percent of Methoxy-XO4 which is non-vascular or vascular in pial (I), cortical (II), or hippocampal (III) tissue (n=3 male J20). **(C)** Normalized percent coverage of Methoxy-XO4 in deep or surface tissue. Coverage is considered the volume of Methoxy-XO4 over the volume of the tissue **(D)** Normalized percent coverage of Methoxy-XO4 in deep tissue or surface vasculature (n=5 male J20). Significance detected through Ordinary One-Way ANOVA. Scale bar = 80µm (AI), 70µm (AII), 30µm (AIII, AVI, AIX) or 100µm (AIV, AV, AVII, AVIII). * = p-value < .05

Methoxy-XO4 has been shown to cross the blood-brain barrier (BBB)(24), and label parenchymal plaques. Studies have shown that Methoxy-XO4 can be administered repeatedly and used in conjunction with *in vivo* two-photon imaging to monitor plaque growth over time(25, 26). Further, >95% of Methoxy-XO4 plaques co-labeled with Aβ antibody 1-day post injection(25, 26). Yet, the exact diffusion volume in live animals is not known. Thus, we utilized premortem Methoxy-XO4 labeling with postmortem 6e10 Aβ antibody co-labeling in a separate cohort of 12-month-old J20 mice. Co-labeling between Methoxy-XO4 and 6e10 Aβ labeling in hippocampal and cortical brain slices were assessed utilizing both an automated image analysis approach in Imaris and manual analysis in ImageJ. Here we identified that ∼70% of Methoxy-XO4 co-labeled with 6e10 stained plaques with no difference in co-labeling observed between surface and deeper brain regions (**Supplemental Fig. 4**). Our results are in line with previously published work demonstrating a high correlation between Methoxy-XO4 labeled plaques and antibody labeled plaques, and demonstrate our approach is suitable for comparing leptomeningeal and parenchymal Ab load.

To determine if the meningeal Aβ load seen across the two AD model systems impacted CBF, laser speckle contrast imaging was employed. This technique evaluates blood flow through the meninges and up to one millimeter of the cortex. Compared to WT controls, J20 and 5XFAD mice exhibited 13.4%, and 10.9% lower CBF, respectively (**Fig. 3A-B, Supplementary Fig 5A-B**). Notably, a significant correlation between meningeal CAA and cerebral blood flow loss was observed in these models at 12 months of age (p = 0.03, **Fig. 3C**). Lateralization and asymmetry in behavior, blood flow, and other pathologies in AD have been debated within the literature. When looking at PET imaging studies, some indicate lateralization(50–52), while others show asymmetry without lateralization(50). Given the scarce research on lateralization in the J20 mouse model of AD, we assessed our animal cohorts and identified no lateralization was noted in blood flow for either J20 or 5XFAD models when compared to age-matched WT littermate controls (**Fig. 3D, Supplementary Fig. 5C**). Pial collateral vessels are critical determinants of patient outcome after an ischemic stroke, as they contain a remarkable adaptive ability to enlarge and compensate for loss of CBF and support retrograde reperfusion to preserve the penumbra. To determine if pial collateral vessels are altered as results of decreased CBF and increased Aβ load in the J20 and 5XFAD mouse models of AD, vessel painting was employed in conjunction with confocal imaging. At 2 months of age, before Aβ accumulation begins, no differences were observed in pial collateral diameter or number in any of the three interconnecting collaterals in J20 mice, when compared to age matched littermate controls (**Supplementary Fig. 6A-D**). However, by 12 months of age when significant meningeal CAA had occurred, the diameter of pial collateral vessels in the J20 MCA-ACA niche were significantly larger compared to control mice (**Fig. 4A-C**). J20 mice displayed significantly fewer collaterals at less than 30μm with greater percentages of collaterals at 31-40μm and 41-50μm in size compared to WT controls (**Fig. 4D**). Interestingly, no differences in collateral diameters were noted in the MCA-PCA and ACA-PCA collateral vessels. Pial collateral vessel tortuosity displayed a similar trend, with significantly greater tortuosity in J20 MCA-ACA pial collaterals compared to WT controls, but no changes in the MCA-PCA and ACA-PCA collaterals (**Fig. 4E**). No changes were seen between genotypes with regard to collateral number, consistent with previous research indicating that collaterals do not form *de novo* (**Fig. 4F**). Taken together, the increased diameter and tortuosity of J20 MCA-ACA collaterals indicate the vessels undergo arteriogenesis during the course of disease progression. Moreover, no significant changes were identified between the sexes, therefore values were combined during analysis (**Supplementary Fig. 7**). No significant alterations were observed in collateral diameter or number in 5XFAD mice at 12 months of age (**Supplementary Fig. 5E-G**). The absence of collateral enlargement in 5xFAD mice indicates that a critical threshold of pial arterial Aβ accumulation is necessary to trigger vascular disruption and arteriogenesis

**Figure 3.**
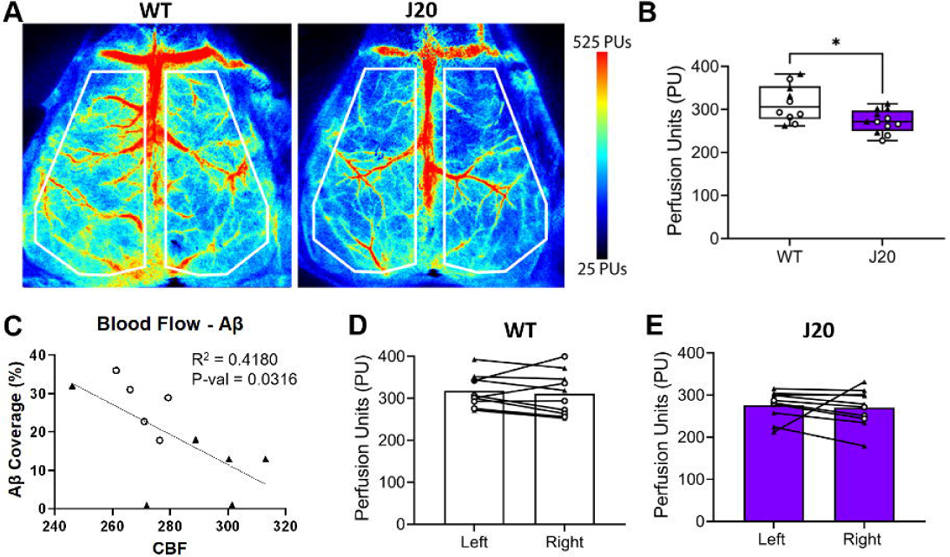
Reduced CBF correlates with pial artery/arteriole vascular network Aβ load in J20 and 5XFAD mice. **(A)** Representative pseudo-colored laser speckle contrast images of WT (n= 6 male, 4 female) and J20 (n= 6 male, 6 female) mice. **(B)** Quantification of averaged hemispheric cerebral blood flow shows a significant reduction in J20 mice compared to WT controls as evaluated by an unpaired t-test. **(C)** Linear regression analysis with 95% confidence bands shows negative correlation between Aβ load and CBF in J20 mice. n= 5 male and 6 female. **** = p-value < .05. Circles = males; Triangles = females; **(D)** No differences were seen in perfusion between the hemispheres of WT (n= 6 male, 4 female) or **(E)** J20 mice (n= 3 males, 6 females) as evaluated by a paired t-test.

**Figure 4.**
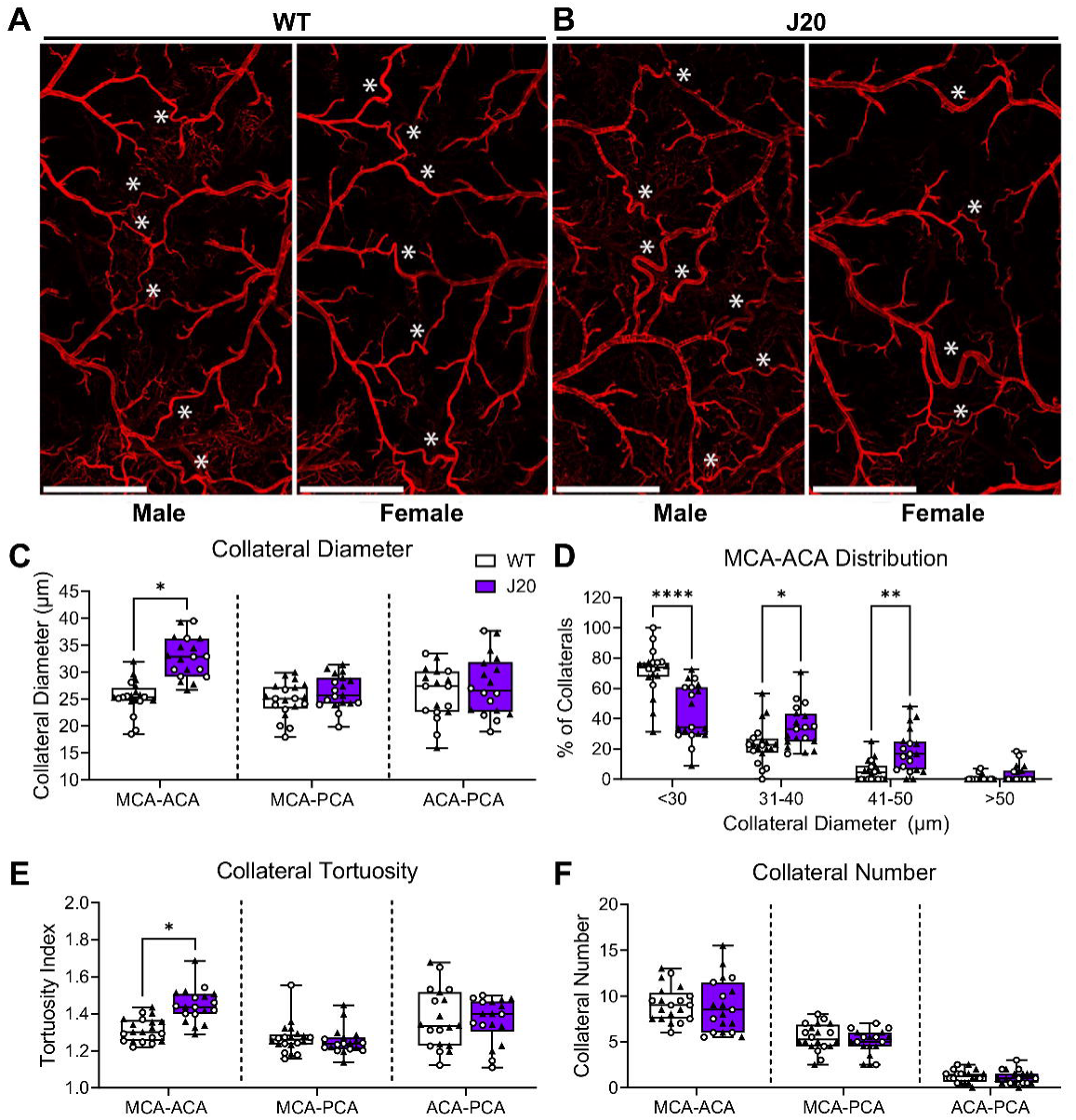
J20 mice exhibit significant alterations in their MCA-ACA collateral niche. Vessel painting was utilized to label the pial artery/arteriole network. **(A)** Representative tiled 4X images of the MCA-ACA pial collateral niche in WT and **(B)** J20 mice. **(C)** Increased collateral diameter was observed in the MCA-ACA niche of J20 mice compared to WT controls. No differences were seen in the MCA-PCA or ACA-PCA collateral niches. **(D)** J20 mice have a significantly higher percentage of MCA-ACA collaterals in the 31-40 and 41-50 um range compared to WT mice. **(E)** Collateral tortuosity was increased only in the MCA-ACA collaterals of J20 mice. **(F)** No significant difference was noted in collateral number between J20 and WT mice in any collateral niche. Significant differences in collateral niches were evaluated using multiple unpaired t-tests and collateral distribution using two-way ANOVA with Sidak multiple comparisons. n= 9 male and 11 female WT, and 8 male and 11 female J20. White * = MCA-ACA collateral blood vessel. Scale bar = 1mm. * = p-value < .05, ** = p-value < .01, **** = p-value < 0.0001. Circles = males; Triangles = Females

## Discussion

In AD, reduced CBF has emerged as a pivotal factor in disease progression, reflecting the intricate interplay between vascular dynamics and neurodegenerative processes. The current study recapitulates the large body of work demonstrating reduced CBF and explores the nuanced relationship between Aβ deposition and reduced CBF. In the J20 model of AD, the most significant accumulation of Aβ was observed on the meningeal arterial vasculature, with male mice carrying a higher burden compared to females. This is the first known report thoroughly characterizing leptomeningeal CAA accumulation, and the first to reveal sex differences in the leptomeninges in an animal model. Deposition of Aβ in the parenchyma of human brains occurs in a hierarchical manner, with Aβ first accumulating in cortical regions, followed by the hippocampus, and then spreading throughout other regions of the brain. Vascular Aβ-deposits occur in the first stage of CAA in leptomeningeal and parenchymal vessels of neocortical regions (53). Our data shows that in the J20 mouse model of AD, Aβ accumulates early and in the highest concentrations first on the leptomeningeal arterial vasculature at 12 months of age when compared to cortical/hippocampal vascular burden and parenchymal burden. Notably, our findings in the J20 mouse model are consistent with studies dating back nearly four decades demonstrating αβ accumulation in the leptomeningeal arterial network early in disease progression (reviewed in (53)). As expected, the level of CAA accumulation in leptomeningeal vasculature correlated with reductions in CBF, suggesting a novel locality for which to target CAA. These findings suggest that age-related meningeal dysfunction, for example, inflammation or perivascular drainage, may initiate surface accumulation of Aβ that drives parenchymal accumulation in a temporospatial manner.

Interestingly, coupled with extensive meningeal CAA, the MCA-ACA leptomeningeal collaterals were enlarged in J20 mice, but no changes were observed in the 5xFAD mouse line. Pial collateral vessels aid in reperfusion which supports rerouting blood flow to vulnerable tissue following ischemic stroke. Pre-existing collateral number and ability to enlarge, through arteriogenesis, is a critical factor in stroke patient outcome. Despite this critical adaptation, limited work has been done to further our understanding of their role in combating the global reduction in blood flow seen in AD models. Work by Zhang et al., found that pial collateral vessel changes seemed to vary by mouse strain(19). In contrast to their results which showed decreases in collateral number and diameter with age in their 5xFAD models evaluated. Conversely, we found that collateral size was enhanced in J20 mice with no alterations to collateral number at 12 months of age. Other findings using 5xFAD mice showed increased collateral diameter from 8-12 months and a significant genotype effect, but the number of vessels decreased with age in both 5xFAD and WT mice (54). The difference in outcome compared to our 5xFAD study may be due to housing conditions or technical approaches used to analyze collateral changes.

Although critical studies examining the pial collateral response in AD are lacking, the connection between CAA CBF is well-established. A recent meta-analysis showed that in patients with AD, 48% displayed moderate to severe CAA. Further, this study showed that in the general population, the prevalence of moderate to severe CAA pathology was 23%(16). While women are more likely to develop CAA in their lifetime, studies have shown that men typically have worse CAA scores, develop CAA earlier, and tend to have worse response to CAA in the form of hemorrhagic disease(55, 56). This is further supported by our work, revealing that male J20 mice have a heavier leptomeningeal CAA burden relative to females. This study underscores the importance of understanding the temporal dynamics of CBF reduction and its relationship to Aβ accumulation in the meningeal arterial vasculature. On its own, a reduction in global CBF is sufficient to cause cognitive dysfunction(57). CAA is known to cause structural changes in the vessel walls, reducing their compliance and impairing CBF regulation, and can reduce the integrity of the blood vessel, increasing risk of hemorrhage. Both factors, in conjunction with the inflammatory response triggered by these Aβ deposits, can lead to exacerbated neuroinflammation and disease progression.

Previously thought to reflect the reduced metabolic demand of advanced AD, it is now well-established that reduced CBF occurs early in AD progression and may contribute to cognitive decline(7, 58), leading to the notion that reduced CBF may serve as an early diagnostic indicator for early AD(4, 7). Reduced CBF may contribute to impaired delivery of oxygen, and glucose, as well as contribute to deficits in clearance of brain metabolic waste and A accumulation(8, 9, 59). Supporting this, in mice overexpressing the Swedish mutation of APP, reduced CBF was observed prior to parenchymal amyloid plaque deposition(10). Reduced CBF correlates with reduced glucose utilization in preclinical AD models(10–12). Vasomotor responses and functional hyperemia appear to be disrupted throughout disease progression. In the early stages, increased neural excitability is linked to an exaggerated hemodynamic response in J20 mice(60). In contrast, in aged APP overexpressing mice (Tg2576), laser speckle imaging correlated vascular Aβ deposition with impaired vasomotor responses to hypercapnia and whisker stimulation-induced hyperaemia(61). Importantly, parenchymal vascular abnormities are described in postmortem AD tissues, including increased tortuosity, thinning of vessel walls and the appearance of wicker-like vascular networks (reviewed in(62)). Similar abnormalities have recently been described using high resolution mapping of hippocampal vasculature in aged (24 month) APP/PS1 mice, including fewer uniform vessels, with a lower mean vascular diameter, more distal fine branching and reduced perfusion areas(63). It is not known if these are early disease events, as these evaluations were performed in late-stage disease. Alternatively, these changes may occur in response to inadequate arterial blood flow because of pial amyloidβ load, as we show, which may potentially cause capillary regression due to a global decrease in flow through micro vessels. Future studies examining the temporal relationship between vascular amyloidβ load in the meningeal arterial network, CBF and parenchymal vascular abnormalities are thus warranted.

Although the mechanisms for CBF impairment in AD are poorly understood, studies have indicated that CBF enhancement may be a viable method of risk reduction for AD which has the potential to reduce CAA and Aβ load. For example, exercise is a robust modifier of AD risk and in mouse models, as long-term access to a running wheel in early or late-stage disease was associated with a decrease in Aβ load and improved microvascular pathology(64–66). Pharmaceutical interventions have also been explored to improve blood flow and reduce cognitive impairment in AD. In humans, a small clinical trial with anti-hypertension drug Nilvadipine showed improved CBF in the hippocampus(67), however a large phase III trial did not find benefits to Nilvadipine treatment in patients with mild to moderate AD(67). Similarly, treatment with anti-hypertension drugs against the receptor for advanced glycation end products (RAGE) failed to show efficacy in a clinical setting and could exacerbate cognitive decline in high doses(68). Therefore, there is a critical need for the development of targeted, safe drugs to restore CBF.

In addition to disrupting normal CBF, CAA is also linked with impaired drainage of waste products from the brains glymphatic system. The glymphatic pathway involves clearance of waste products, including Aβ, through perivascular spaces, extending into the meninges, before reaching lymphatic vessels. Recent work has suggested that clogging or dysfunction of the glymphatic system could potentially impair Aβ clearance, contributing to CAA development. Impaired meningeal lymphatic drainage exacerbates the microglial inflammatory response in AD suggesting that enhancement of meningeal lymphatic function combined with immunotherapies could lead to better clinical outcomes(69).

### Limitations and future directions

There exist inherent limitations of LSCI in capturing deep brain structures, with reported penetration depths ranging from 300 µm to 1mm(70, 71) and LSCI does not provide absolute quantification of CBF. Yet, LSCI offers valuable insights into superficial cortical perfusion dynamics. Here we emphasize its utility in quantifying superficial cortical and meningeal perfusion, where alterations in blood flow can have significant implications for neurovascular function. LSCI also enables real-time, non-invasive assessment of microvascular perfusion, providing a practical means of monitoring changes in CBF.

Isoflurane, used here, is a potent vasodilator that increases basal CBF and can influence arteriole reactivity(72, 73). Anesthetic choice is known to impact CBF measurements. In a separate set of experiments, we identified that the differences in CBF were maintained between WT and J20 mice using ketamine/xylazine as the anesthetic agent (**Supplemental Figure 8**). It should be noted that overall CBF is lower using ketamine/xylazine which is associated with lower cerebral partial pressure of oxygen and moderate hypoxia(74, 75). Future studies evaluating other anesthetic agents are warranted.

Finally, we did not test directionality of blood flow within the collaterals to investigate whether collateral remodeling could directly aid in redistribution of CBF. Yet, previous studies have shown that collateral vessel enlargement and remodeling improve collateral flow capacity, facilitating reperfusion in occlusion scenarios(29, 33, 76, 77). Our findings of increased collateral diameter and tortuosity in AD mice align with this concept of adaptive remodeling that may be initiated to compensate for changes in CBF.

### Conclusions

Our study underscores the pivotal role of CBF in AD, shedding light on the intricate interplay between vascular dynamics and CAA. The observed enhancement of collateral size in J20 mice, coupled with increased meningeal CAA, prompts consideration of collateral vessels as potential therapeutic targets for AD. Overall, our work contributes to the evolving narrative on AD vascular dysfunction and calls for continued exploration of novel therapeutic avenues.

## Declarations

### Ethics Approval and Consent to Participate

This study was approved under the Institutional Animal Care and Use Committee at Virginia Tech under protocol number 21-079 and 24-037

### Consent for Publication

Not applicable

### Availability of Data and Materials

The datasets used and/or analyzed during the current study are available from the corresponding author on reasonable request.

### Author’s Contributions

A.M.K. was responsible for design, execution, analysis, and interpretation of laser speckle contrast imaging, vessel painting, imaging, collateral morphology analysis, writing and editing. J.L.B. was responsible for design, execution, and interpretation of Methoxy-X04 injections, tissue clearing, IHC, imaging, CAA burden analyses, writing, editing, and communication. J.L. was responsible for design and execution of animal husbandry, genotyping, Methoxy-X04 injections, tissue clearing, imaging, and writing. Y.P. and S.W. were responsible for CAA burden analysis assistance. H.S. carried out project supervision and funding acquisition along with M.L.O. M.H.T. and M.L.O. conceived the idea for the study and carried out project supervision, data interpretation and manuscript writing.

## Acknowledgments

Not applicable

## Abbreviations

5XFAD: APPSwFlLon,PSEN1*M146L*L286V)6799Vas/Mmjax
Aβ: Amyloid beta
ACA: Anterior cerebral artery
AD: Alzheimer’s disease
BBB: Blood brain barrier
CAA: Cerebral amyloid angiopathy
CBF: Cerebral blood flow
CNS: Central nervous system
J20: B6J.Cg-*Zbtb20^Tg(PDGFB-APPSwInd)20Lms^*/Mmjax, RRID:MMRRC_034836-JAX
LSCI: Laser speckle contrast imaging
MCA: Medial cerebral artery
MCI: Mild cognitive impairment
PBS: Phosphate buffered saline
PCA: Posterior cerebral artery
THF: Tetrahydrofuran
WT: Wildtype

## Supplemental Figure Legends

**Supplemental Figure 1. Collateral identification from vessel painted images. (A)** Representative tiled 4X confocal image of vessel painted male J20 brain. Collateral blood vessels are identified within the dashed box by their characteristic tortuosity and typically non-physiological, wide angles of attachment to the parent arteries. These intercollateral vessels, which connect two independent arteries, are positioned directly between the distal-most arterioles of the parent arteries. **(B)** Angle of attachment to the parent artery of collateral vessels. These vessels often exhibit wide, variable angles that can impact blood flow dynamics. **(C)** Collateral tortuosity is calculated using the straight line of collateral length (white) and freehand length (dashed yellow). The formula used is: tortuosity = freehand length/straight-line length X 100. Scale bars = 1mm for panel A and 500um for panels **B** and **C**.

**Supplemental Figure 2. Comparison of FIJI (ImageJ) and Imaris A**β **coverage analyses in 12-month-old J20 mice. (A)** Average coverage of entire leptomeningeal vasculature by Aβ as quantified by ImageJ and Imaris analysis with **(B)** matched samples. No significance was detected. Significance tested for using unpaired (A) and paired (B) t-test. N = 7 female J20 and 2 male J20. Circles = males; triangles = females.

**Supplemental Figure 3. Leptomeningeal Aβ coverage in the 12 month old male and female WT and 5XFAD Mouse (A**) Representative images of vessel paint (red) and Methoxy-XO4 (cyan) in 12-month-old WT and 5X FAD mice and corresponding 3D Imaris reconstructions (scale bar = 500µm) **(B)** Representative zoomed image of MCA-ACA collateral bed in J20 animal (Anterior Cerebral Artery (ACA) = green, collaterals = orange, Medial Cerebral Artery (MCA) = red, distal arterials = gold, scale bar = 200µm) **(C)** % Aβ coverage of the MCA, MCA-ACA Collaterals, and ACA with significance tested by RM One-Way ANOVA test. N = 4 male 5X FAD, 2 female 5X FAD, 4 male WT, and 5 female WT. Circles = males; triangles = females

**Supplemental Figure 4. Methoxy XO4 and antibody labeling approaches to identify parenchymal plaques. (A)** Representative images of Methoxy-XO4 (blue), GFAP (green) and 6e10 (red) in cortex and hippocampus of male J20 mice. White arrows mark plaques which are co-labeled with Methoxy-XO4 and 6e10 (scale = 50µm) **(B)** The number of Methoxy-XO4 labeled plaques within 10µm of GFAP signal (II) The number of Methoxy-XO4 labeled plaques in contact with 6e10 signal (III) Methoxy-XO4 plaque size and (IV) number and (V) percentage of Methoxy-XO4 plaques that colabel with 6e10 and 6e10 plaques that colabel with Methoxy-XO4 in cortex and hippocampus. Significance was tested through paired t-test.

**Supplemental Figure 5. Despite reduced blood flow, collateral diameter and collateral number is unaltered in aged FXFAD mice. (A)** Pseudocolored laser speckle contrast images of WT (n= 6 male, 8 female) and 5xFAD (n= 5 male, 6 female) mice at 12 months of age. **(B)** Analysis by unpaired t-test showed significant reduction in cerebral blood flow in 5xFAD mice compared to WT littermates. **(C)** No differences were seen in hemispheric perfusion **(D)** No significant differences were noted in cerebral blood flow between 5xFAD and J20 mice, or their control littermates. **(E)** Representative confocal images of the MCA-ACA pial collateral niche of WT and 5xFAD mice. **(F)** No differences were found in collateral diameter or **(G)** number between 5xFAD (n = 4 male and 5 female) and WT mice (n = 4 male and 5 female). Significant differences in collateral niches were evaluated using multiple unpaired t-tests. Circles = males; Triangles = females

**Supplemental Figure 6. Early life J20 mice do not show changes in collateral remodeling. (A)** Representative vessels painted and Methoxy-XO4 images of WT and **(B)** J20 mice at 2 months of age, prior to Aβ accumulation. **(C)** Collateral diameter and **(D)** number is unaltered in any of the intercollateral niches between WT (n = 5 female) and J20 (n = 5 female) mice. Significant differences in collateral niches were evaluated using multiple unpaired t-tests. White * = MCA-ACA collateral blood vessel. Scale bar = 1mm. Significant differences in collateral niches were evaluated using multiple unpaired t-tests. Circles = males; Triangles = Females

**Supplemental Figure 7. Collateral characteristics do not differ between sexes. (A)** No significant sex differences were noted in each genotype for collateral diameter, **(B)** number, **(C)** or tortuosity. Significant differences in collateral niches were evaluated using multiple unpaired t-tests. n= 9 male and 11 female WT, and 8 male and 11 female J20. Circles = males; Triangles = Females

**Supplemental Figure 8. Effect of anesthetic choice on cerebral perfusion measured by laser speckle contrast imaging. (A)** Representative pseudocolored laser speckle contrast images of WT and J20 mice anesthetized with either isoflurane or ketamine/xylazine. **(B)** Quantitative analysis of averaged hemispheric perfusion demonstrates significantly reduced cerebral perfusion in J20 mice anesthetized with ketamine/xylazine. Statistical analysis was performed using an unpaired t-test. *p < 0.05, N = 3–5 males.

